# Activation of the fear-responsive anterior hypothalamic area promotes avoidance and triggers compulsive grooming behavior in mice

**DOI:** 10.1101/2022.09.06.506804

**Authors:** Brenton T. Laing, Megan S. Anderson, Aishwarya Jayan, Anika S. Park, Lydia J. Erbaugh, Oscar Solis, Danielle J. Wilson, Michael Michaelides, Yeka Aponte

**Affiliations:** Neuronal Circuits and Behavior Section, National Institute on Drug Abuse Intramural Research Program, National Institutes of Health, Baltimore, MD 21224-6823, U.S.A.; Biobehavioral Imaging and Molecular Neuropsychopharmacology Unit, National Institute on Drug Abuse Intramural Research Program, National Institutes of Health, Baltimore, MD 21224-6823, U.S.A.; Department of Psychiatry and Behavioral Sciences, Johns Hopkins University School of Medicine, Baltimore, MD 21205, U.S.A.; The Solomon H. Snyder Department of Neuroscience, Johns Hopkins University School of Medicine, Baltimore, MD 21205, U.S.A.

## Abstract

The anterior hypothalamic area (AHA) is a key brain region for orchestrating defensive behaviors. Here, we first examined AHA activity patterns during fear conditioning using *in vivo* functional imaging. We observed that neuronal activity in the AHA increases during both foot shock delivery and foot-shock associated auditory cues. Moreover, we used a combination of optogenetics and behavioral assays to determine the functional connectivity between the ventromedial hypothalamus (VMH) and the AHA. We found that photoactivation of the VMH→AHA pathway is aversive and triggers compulsive grooming behavior. Furthermore, we observed spatial and temporal changes of grooming behavior during the periods following VMH→AHA photoactivation. Interestingly, whole brain metabolic mapping using positron emission tomography (PET) combined with optogenetic activation of the VMH→AHA pathway in anesthetized mice revealed the amygdala as a downstream area activated by the stimulation of this pathway. Together, our findings show that the AHA responds to threat and that such increases in activity are sufficient to trigger compulsive grooming behavior. Thus, our results may help to understand some neuropsychiatric disorders characterized by repetitive and compulsive behaviors.

## INTRODUCTION

Epidemiological data indicate that post-traumatic stress disorder (PTSD) and obsessive-compulsive disorder (OCD) share strong comorbidity^1, 2^ but the extent to which such interactions are influenced by common brain circuit mechanisms is not well understood. Association of salient cues with threatening stimuli aids in prediction of environments that may present threat^3-5^.Previous studies have demonstrated that the ventromedial hypothalamus (VMH) and the anterior hypothalamic area (AHA) are critical to this process. Robust escape behaviors such as increases in locomotor activity and jumping were triggered by the activation of the VMH→AHA pathway^6^. Moreover, activation of hippocampal inputs to the AHA promotes contextual control of such behaviors^7^. Thus, these findings raise questions about the etiological role of AHA neuronal activation in stress-related and compulsive disorders^8^.

While it is known that AHA circuitry is involved in escape behaviors, it remains largely unstudied whether the AHA only encodes for information related to innate threats or also plays a generalized role in learning. Moreover, whether the activation of the AHA threat response circuitry causes changes that persist beyond stimulation or if it simply acts as an “on-and-off switch” have yet to be determined. Previous studies showed that repeated activation of other fear-associated circuits leads to persistent compulsive grooming behavior^9, 10^. Therefore, we also sought to investigate changes in activity patterns of key threat regulatory regions triggered by the photoactivation of AHA circuitry.

Here, we addressed these questions by using a combination of *in vivo* functional imaging, optogenetics, and behavioral assays. First, the genetically encoded calcium indicator GCaMP and a detachable miniscope were used to record AHA neuronal activity in freely moving mice^11^ during a classic foot shock-cue conditioning paradigm^12^. Moreover, we analyzed AHA neuronal activity changes during foot shock delivery and foot-shock associated auditory cues. Second, we used optogenetics to photoactivate the VMH→AHA pathway and recorded both the spatial and temporal behavioral outputs triggered by the activation of this pathway. Finally, using positron emission tomography (PET) and optogenetics, we observed that the amygdala is a downstream area activated by the stimulation of VMH→AHA pathway.

## MATERIALS AND METHODS

### Animals

All experimental protocols were conducted in accordance with U.S. National Institutes of Health Guidelines for the Care and Use of Laboratory Animals and with the approval of the National Institute on Drug Abuse Care and Use Committee. Male and female wild type (WT) mice (C57BL6/J background; IMSR Cat# JAX:008069, RRID:IMSR_JAX:008069; The Jackson Laboratory, ME, U.S.A.) were used. Prior to surgery, mice were group housed with littermates in temperature and humidity-controlled rooms on a 12 h light/dark cycle with *ad libitum* access to water and rodent chow (PicoLab Rodent Diet 20, 5053 tablet, LabDiet/Land O’Lakes Inc., MO, U.S.A.). As a function of breeding output, groups were approximately age and sex matched prior to surgery.

### Stereotaxic viral injections

Stereotaxic microinjections were performed using a stereotaxic apparatus (David Kopf Instruments, Tujunga, CA, U.S.A.) and micromanipulator (Narishige International U.S.A. Inc, Amityville, NY, U.S.A.) with custom-pulled glass pipettes (20−30 μM tip diameter). Mice were anesthetized using isoflurane (4% for induction and 1.5 – 2% for surgery) and administered post-operative ketoprofen (5 mg/kg, s.c.) for analgesia. For *in vivo* functional imaging experiments, an adeno-associated virus (AAV) was injected into the anterior hypothalamic area (AHA; 4x50 nl for 200 nl total per mouse; AP: −0.70 and −0.94, ML: +0.425, DV: −5.50 and −5.30) for the expression of the genetically encoded calcium indicator GCaMP^11^ and a GRIN lens (500-μm-diameter; Snap-in Imaging Cannula Model L-V, Doric Lenses Inc., Québec, QC, Canada) was implanted to record AHA neuronal activity (AP: −0.85, ML: +0.45, DV: −5.22). For behavioral and PET scan experiments involving photoactivation of the VMH→AHA pathway, viral injections into the VMH (AP: −1.7, ML: +0.3, DV: −5.75) were performed and custom fabricated optical fibers^13^ (200-μm-core, 0.48 NA, 4.8-mm-length) were unilaterally implanted dorsal to the AHA (AP: −0.85, ML: +0.45, DV: −4.8). For optogenetic experiments, no control mice were excluded based on optimal viral transduction. However, channelrhodopsin (ChR2)-expressing mice were excluded if predominant viral expression was not medial to the fornix and targeted at the ventral half of the third ventricle. Therefore, we excluded *n* = 4 ChR2-expresing mice that did not meet such criteria. A total of *n* = 11 mice in the control group and *n* = 7 ChR2-expressing mice in the stimulation group were used for experiments.

Viruses used include: (1) rAAV2/1-Camk2a-hChR2(H134R)-EYFP-WPRE, titer: 5.0 × 10^12^ GC/ml (Addgene viral prep # 26969-AAV1, RRID:Addgene_26969; Addgene, MA, U.S.A.)^14^, (2) rAAV2/1-Camk2a-eYFP-WPRE, titer: 5.0 × 10^12^ GC/ml (Addgene viral prep # 105622-AAV1, RRID:Addgene_105622), (3) rAAV2/9-Syn-jGCaMP7s-WPRE, titer: 5.0 × 10^13^ GC/ml (Addgene viral prep # 104487-AAV1, RRID:Addgene_27056)^15^.

### Foot shock conditioning during functional imaging

Functional imaging in freely moving mice was performed by recording jGCaMP7s fluorescence from AHA neurons with a Doric Lenses, Inc. microendoscope interfaced with an implanted GRIN lens^16^ using Doric Neuroscience Studio software v5.1 (RRID:SCR_018569). Approximately 4 weeks after surgery, mice were habituated to experimenter handling for 5 minutes daily for one week.

Experimental procedures were conducted approximately 5 weeks post-surgery. This included one day of foot shock conditioning and one day of cue-response testing^12^. Each day consisted of a 5-min test. The protocol for day 1 (i.e., 120 s wait, 10 s tone, 5 s foot shock, 120 s wait, 10 s tone, 5 s foot shock, 35 s extra recording) was similar to the protocol for day 2 (i.e., 120 s wait, 10 s tone, 125 s wait, 10 s tone, 10 s, 30 s extra recording) except the foot shock was omitted on day 2. Acquisition parameters were set to 100-ms exposure and 0 gain. Illumination power remained constant for both sessions. Fluorescence signals were extracted and converted into normalized intensity values as Z-score using the miniscope pipeline^17^. Freezing behavior was identified by using ANY-maze video tracking software v6 (RRID:SCR_014289; Stoelting Co., IL, U.S.A.) with default settings and 1-s detection time. Custom MATLAB scripts were used for perievent alignment to protocol and behavior time stamps (MATLAB R2020a, RRID:SCR_001622; MathWorks, Natick, MA, U.S.A.).

### Optogenetic manipulations and behavioral assays

All behavioral tests were conducted within the light phase of the light/dark cycle. Mice were acclimated to the testing room for at least 1 h prior to each experiment. Before testing, mice were tethered to a 450-nm laser (Doric Lenses, Inc.) via fiber-optic patch cords. The photostimulation protocols (10-ms pulse width at 20 Hz) were generated using Doric Neuroscience Studio software v5.1. Videos were manually scored for jumping, grooming, and rearing behaviors with ANY-maze software v6 by an observer blinded to treatment groups. Notably, grooming was defined as licking, rubbing, scratching, or nibbling at any part of the body.

### Open field test (OFT)

To detect the effects triggered by photoactivation of the VMH→AHA pathway, mice were placed in open field arenas (30 cm x 30 cm) with a thin layer of mice bedding on the arena floor. Arenas were placed inside isolation chambers illuminated to approximately 150 lx. The 18-min testing session was divided into six alternating 3-min epochs. This bin length has been demonstrated to modulate amygdala-dependent behavior^18^. During the first, third, and fifth epochs, the laser was off. During the second, fourth, and sixth epochs the laser was on. An overhead camera and ANY-maze software v6 were used to record and assess mice locomotion and location.

### Real-time place preference (RTPP)

For real-time place preference experiments, a standard-sized rat cage (20 cm × 40 cm) with black opaque walls and a layer of bedding was placed in an isolation chamber with the overhead lights turned off. Laser stimulation was paired with one side of the chamber (laser-ON side) and was consistent across sessions. Mice were placed in the laser-OFF side and could freely transition between the two sides for 20 min. Average speed and total time spent in each side of the chamber were calculated by ANY-maze software v6. Immobility was defined as the absence of movement in the X, Y, and Z space^6^. For immobility detection, the sensitivity slider was set to 80% with a minimum duration threshold of 2 s. Grooming was manually scored offline.

### Persistent stimulation

Extending the duration of optogenetic stimulation can trigger changes that are different from those during short duration stimulation^19^. To measure the effects of extending the time course of VMH→AHA photoactivation, mice were placed in a testing apparatus for 3 h per day over three consecutive days (Days 1−3). During these sessions, mice were tethered to a laser and received light pulses (10 ms, 20 Hz, 10−15 mW) for 3 h. An additional 3 min of video was acquired each day after shutting off the lasers to obtain post-session grooming scores. Data analysis was performed in MATLAB (R2020a, RRID:SCR_001622; MathWorks, Natick, MA, U.S.A.) to extract animal velocity 5 s before, during, and after photostimulation.

### Positron emission tomography (PET)

Mice from behavioral experiments were used for PET. As described above, mice were injected with AAV-Camk2a-ChR2:YFP or AAV-Camk2a-YFP into the VMH and an optical fiber was implanted dorsal to the AHA in the right hemisphere. Mice were fasted for ∼16 h before the experiment. On the day of the experiment, mice were anesthetized with 2% isoflurane and placed on a custom-made bed in a nanoScan small animal PET/computed tomography (CT) scanner (Mediso Medical Imaging Systems, Budapest, Hungary). Mice were then injected (i.p.) with 13 MBq of 2-deoxy-2-[18F]fluoro-D-glucose (FDG; Cardinal Health, Inc., Dublin, OH, U.S.A.) and scanned for 30 min using a dynamic acquisition protocol followed by a CT scan. During FDG uptake, mice were photostimulated using 3-min OFF/ON blocks (10−15 mW, 10-ms pulse, 20 Hz). PET data were reconstructed and corrected for dead-time and radioactive decay^20^. All qualitative and quantitative assessments of PET images were performed using the PMOD software environment (RRID:SCR_016547; PMOD Technologies LLC, Zurich, Switzerland) and Mediso’s Nucline software. The data were reconstructed in frames corresponding to the blocks of stimulation and the dynamic PET images were co-registered to magnetic resonance imaging (MRI) templates using PMOD’s built-in atlases. All statistical parametric mapping analyses were performed using MATLAB R2016 (RRID:SCR_001622) and SPM12 (RRID:SCR_007037; https://www.fil.ion.ucl.ac.uk/spm/software/spm12/; University College London, London, U.K.) and evaluated at the *p* < 0.05 level using the Probabilistic Threshold-Free Cluster-Enhancement (pTFCE) method and multiple comparison correction^21^ and a cluster extent correction threshold of 100 contiguous voxels (k = 100).

### Histology

Mice were deeply anesthetized with isoflurane and transcardially perfused with 1x phosphate buffered saline (PBS) followed by 4% paraformaldehyde (PFA) in 1x PBS. Whole brains were removed and post-fixed in 4% PFA at 4 °C for at least 24 h before further processing. For all experiments, tissue was embedded in 4% agarose in PBS and 50-μm free-floating, coronal brain sections were collected using a vibratome (Leica VT1200S vibrating microtome, RRID:SCR_020243; Leica Biosystems Inc., IL, U.S.A.), mounted with DAPI-Fluoromount-G aqueous mounting medium (Cat # 17984-24, Electron Microscopy Sciences, PA, U.S.A.) onto Superfrost Plus glass slides (Cat # 48311-703, VWR International, PA, U.S.A.), and imaged with an AxioZoom.V16 fluorescence stereo zoom microscope (Carl Zeiss Microscopy, NY, U.S.A.) using Zen 2012 software (Carl Zeiss Microscopy LLC).

For representative images of axon fields in the AHA, sections were immunostained for GFP using chicken anti-GFP polyclonal antibody (1:1000, Cat # GFP-1020, RRID:AB_10000240, Aves Labs, Inc., OR, U.S.A.). Slices were washed in PBS 6 × 10 min each. Then, slices were blocked for 1 h in PBS + 0.3% Triton X-100 + 3% normal donkey serum. Samples were incubated overnight at room temperature in primary antibody diluted in block solution. The next day, slices were washed 6 × 10 min each followed by incubation in secondary antibody in block solution: goat anti-chicken Alexa Fluor 488 (1:500, Cat # A11039, RRID:AB_2534096; Thermo Fisher Scientific, MA, U.S.A.). Slices were then mounted onto Superfrost Plus glass slides (VWR International) and imaged with an Axiozoom.V16 fluorescence stereo zoom microscope using Zen 2012 software (Carl Zeiss Microscopy LLC) or Keyence BZ-X710 (Keyence Corp., Itasca IL, U.S.A.).

### Experimental design and statistical analysis

Data are reported as mean ± s.e.m. GraphPad Prism 8 software (RRID:SCR_002798; GraphPad, La Jolla, CA, U.S.A.) was used for all graphs and statistical analyses. See **Supplementary Table 1** for specific information about each test. Generally, Student’s *t*-test was used to determine differences between groups when there were not repeated measures. QQ plots and residual plots were assessed for normality. For Student’s *t*-tests, Welch’s correction was applied when variance was different between groups. Two-way repeated measures ANOVA was used to detect differences between groups and within groups when appropriate. Sphericity was not assumed and Greenhouse-Geiser corrections were made for all experiments. Sidak’s multiple comparisons tests were used for further evaluation when significant main effects were detected. For multiple comparisons of foot shock testing experiments, only comparison against the pre-test baseline was conducted. The number of experimental units is depicted as ‘*n*’ for number of mice for each experiment, except for the functional imaging data where ‘*n*’ is the number of cells.

## RESULTS

### Foot-shock conditioned auditory cues evoke an increase in AHA neuronal activity

A recent study showed that contextual learning influences AHA neuronal activity in part through hippocampal inputs that enhance goal-directed escape behavior^7^. Therefore, we first investigated whether a classic foot shock-cue conditioning paradigm modulates AHA neuronal activity. For this, we specifically targeted an adeno-associated virus (AAV) expressing jGCaMP7s to AHA neurons and visualized their activity using a miniscope attached to an implanted GRIN lens (**Figure 1A**) during shock-paired conditioning (**Figure 1B**). We observed that shock-paired auditory cues evoke a significant increase in immobility or freezing behavior (**Figure 1C**). Next, we analyzed changes in fluorescence (**Figure 1D**) off-line using a miniscope analysis pipeline to identify spatial footprints for individual cells (**Figure 1E**). The activity of individual AHA neurons and mice locomotor speed were aligned with the tone and foot shock events (**Figure 1F**) revealing that the auditory tone did not elicit changes in AHA neuronal activity or speed while the foot shock delivery evoked a significant increase in both activity and locomotion on Day 1 (**Figure 1G, I, J**). On Day 2, after shock-paired conditioning, the tone itself triggered a significant increase in AHA neuronal activity and locomotion (**Figure 1H, I, J**) suggesting that AHA neurons are modulated by fear conditioning. Moreover, after shock-paired conditioning, AHA neuronal activity was significantly lower during the shock and post-shock periods established on Day 1 (**Figure 1I**).

**Figure 1.**
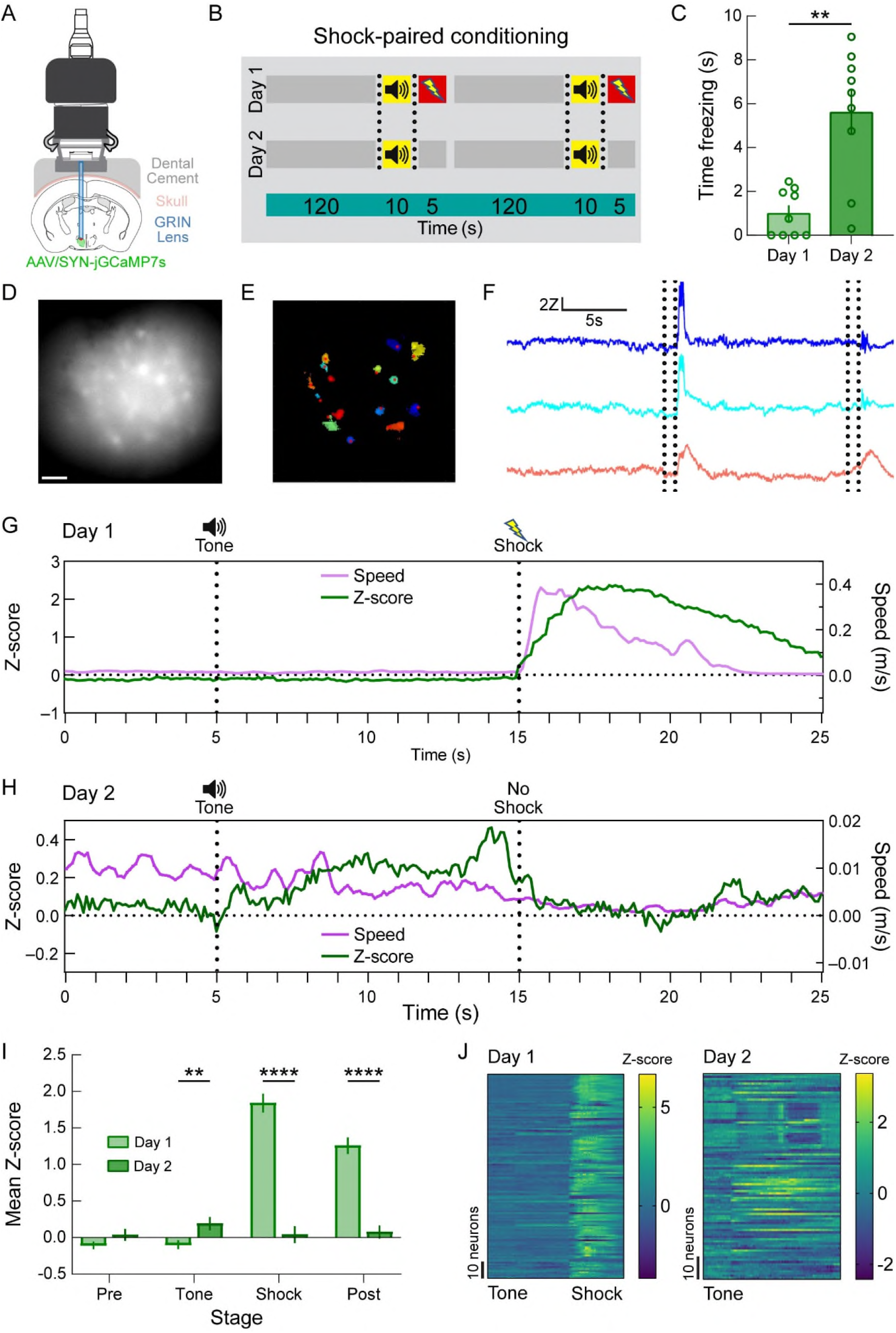
Changes in AHA neuronal activity during foot shock-cue conditioning. **(A)** Viral microinjection strategy for expression of jGCaMP7s with GRIN lens implantation for imaging calcium-dependent signal in the AHA. **(B)** Experimental timeline depicting the shock-paired conditioning session on Day 1 and testing session on Day 2 with two trials for each session. **(C)** Significant differences in two-trial average of freezing behavior are detected after pairing the cue with the shock. **(D)** Representative image of *in vivo* AHA:GCaMP expression. Scale bar = 500 μm. **(E)** Randomly colored spatial footprints of individual neurons used for signal extraction. **(F)** Representative Z-scored traces from individual neurons during a Day 1 session. For each pair of black dotted lines, the first line indicates the time of tone onset, and the second line indicates the time of shock delivery. **(G)** Two-trial average of Z-scored calcium signal intensity (green, *n* = 110 cells) and locomotor speed (purple, *n* = 8 mice; 4 male and 4 female) demonstrating significant increases in AHA activity and speed during shock delivery. No effects are detected during tone presentation on Day 1. **(H)** Two-trial average of Z-scored calcium signal intensity (green, *n* = 90 cells) and locomotor speed (purple, *n* = 8 mice; 4 male and 4 female) demonstrating increases in AHA activity during the tone on Day 2. **(I)** Binned data show a significant increase in Z-scored fluorescence intensity during the tone on Day 2 and a significantly lower Z-score in the absence of shock. **(J)** Heat maps showing changes in AHA neuronal activity for each neuron on Days 1 and 2.

We next examined the activity patterns of AHA neurons from mice that did (*n* = 5/8) and did not (*n* = 3/8) exhibit freezing behavior after shock-paired conditioning (**Supplementary Figure 1A**). We observed that AHA neuronal activity was significantly lower in mice that did not freeze during the tone on Day 2 compared to mice that did (**Supplementary Figure 1B−D**). Furthermore, machine learning models used to classify the mice into freeze and non-freeze groups revealed that substantial information about behavioral selection resides within the AHA. Cubic support vector machine (SVM), linear SVM, quad SVM, weighted k nearest neighbors (KNN), cosine KNN, and fine KNN generated acceptable precision scores (**Supplementary Figure 1E**). Notably, linear SVM produced very poor recall scores, indicating multidimensionality of the data. Quad SVM, weighted KNN, and cosine KNN produced near-chance level recall scores (**Supplementary Figure 1F**). However, fine KNN and cubic SVM performed quite well for this analysis, indicating detection of distinct clusters of neurons predictive of freezing behavior. The approximately equal precision performance for all models combined with the strength of fine KNN and cubic SVM for recall leads to generation of the highest F1 scores in those models (**Supplementary Figure 1G**).

### Photoactivation of the VMH→AHA pathway triggers escape followed by grooming behavior

Previous studies showed that activation of the VMH→AHA pathway promotes escape behaviors^6^. However, the repertoire of behavioral outputs triggered by the activation of this pathway has yet to be determined. For this, we targeted the VMH→AHA pathway for optogenetic and behavioral manipulations (**Figure 2A, B**). We unilaterally injected a viral vector driving the expression of either YFP (control fluorophore; **Figure 2C**; left panel) or channelrhodopsin-2 (ChR2:YFP; light-sensitive neuronal activator; **Figure 2D**; left panel) under the control of the Camk2a promoter to activate glutamatergic neurons in the VMH of wild type mice and implanted an optical fiber above the VMH axonal projections in the AHA (**Figure 2C, D**; right panels). Next, we placed the mice on an open field apparatus^22^ and photostimulated the VMH→AHA pathway using a fixed interval schedule for the delivery of light pulses (i.e., 3-min alternating periods between “ON” and “OFF”). Consistent with previous studies^6^, we observed a significant effect on locomotor distance (**Figure 2E**) and jumping behavior (**Figure 2F**). Moreover, a significant increase in time spent immobile was detected when photostimulation was OFF (**Figure 2G**). We found significant increases in the time spent grooming during the OFF epochs that followed photostimulation (**Figure 2H**), and we observed a significant suppression of rearing behavior (**Figure 2I**) that persisted between photostimulation epochs. There was no significant effect on time in center zone (**Figure 2J**). Together, these results indicate that the characteristic escape behavior evoked by VMH→AHA activation is followed by the emergence of grooming behavior.

**Figure 2.**
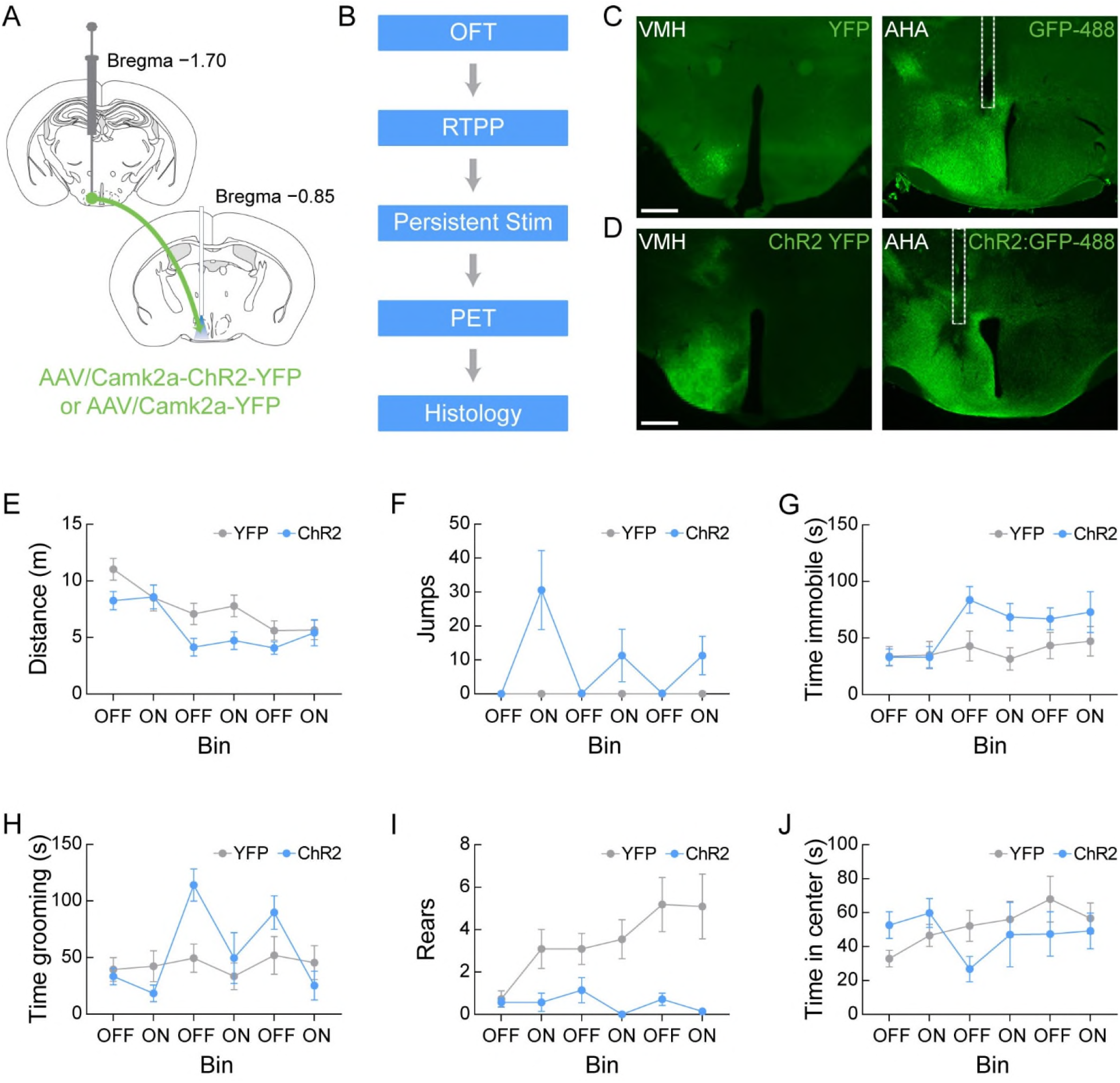
Photoactivation of the VMH→AHA pathway increases jumping and grooming behaviors. **(A)** Viral microinjection strategy for the expression of ChR2:YFP or YFP control fluorophore in VMH glutamatergic neurons with an optical fiber implanted above the VMH axonal projections. **(B)** Timeline for optogenetic behavioral experiments. **(C)** Representative image depicting YFP fluorophore control expression in the VMH (left) and the immunostained (GFP-488) axon field with fiber optic tract (dotted white outline) in the AHA (right). **(D)** Representative image depicting ChR2:YFP expression in the VMH (left) and the immunostained (GFP-488) axon field with fiber optic tract (dotted white outline) in the AHA (right). **(E)** No significant between groups differences are detected for locomotor behavior in the OFT. **(F)** A significant increase of jumping behavior is observed during photostimulation of the ChR2-expressing mice compared to periods without stimulation and fluorophore controls. **(G)** A significant increase in time spent immobile emerges in ChR2-expressing mice after stimulation compared to YFP control mice. **(H)** A significant increase in time spent grooming is observed in ChR2-expressing mice after photostimulation. **(I)** A significant suppression of rearing behavior occurs in ChR2-expressing mice compared to fluorophore controls. **(J)** No effects on time spent in the center zone are detected. See **Supplementary Table 1** for corresponding statistics.

### Photoactivation of the VMH→AHA pathway is aversive and triggers compulsive grooming behavior

We next used a real-time place preference paradigm (RTPP)^23^ to examine the effects of VMH→AHA photostimulation on place preference or aversion. For this test, photostimulation was paired with one side of the arena (Laser-ON zone) and the laser remained off on the other side of the arena (Laser-OFF zone). Mice were permitted to freely move around the arena during testing. In congruence with previous indications that activation of the VMH→AHA pathway triggers avoidance behavior, we observed a significant reduction in the amount of time the ChR2-expressing mice spent in the Laser-ON zone of the arena compared to YFP control mice (**Figure 3A, B**). This was further indicated by a significant reduction in the average duration in the Laser-ON zone (**Figure 3C**). However, there was not a significant reduction of entries to the Laser-ON zone (**Figure 3D**), suggesting a potential lack of behavioral learning. Additionally, the ChR2-expressing mice spent significantly more time immobile (**Figure 3E**). Furthermore, even when normalized for the amount of time in the Laser-OFF zone, the ChR2-expressing mice exhibited significantly higher time spent grooming than the control mice (**Figure 3F**). Moreover, a significant suppression of digging (**Figure 3G**) and rearing (**Figure 3H**) was observed in the ChR2-expressing mice compared to controls. Together, these results further demonstrate that activation of the VMH→AHA pathway is aversive.

**Figure 3.**
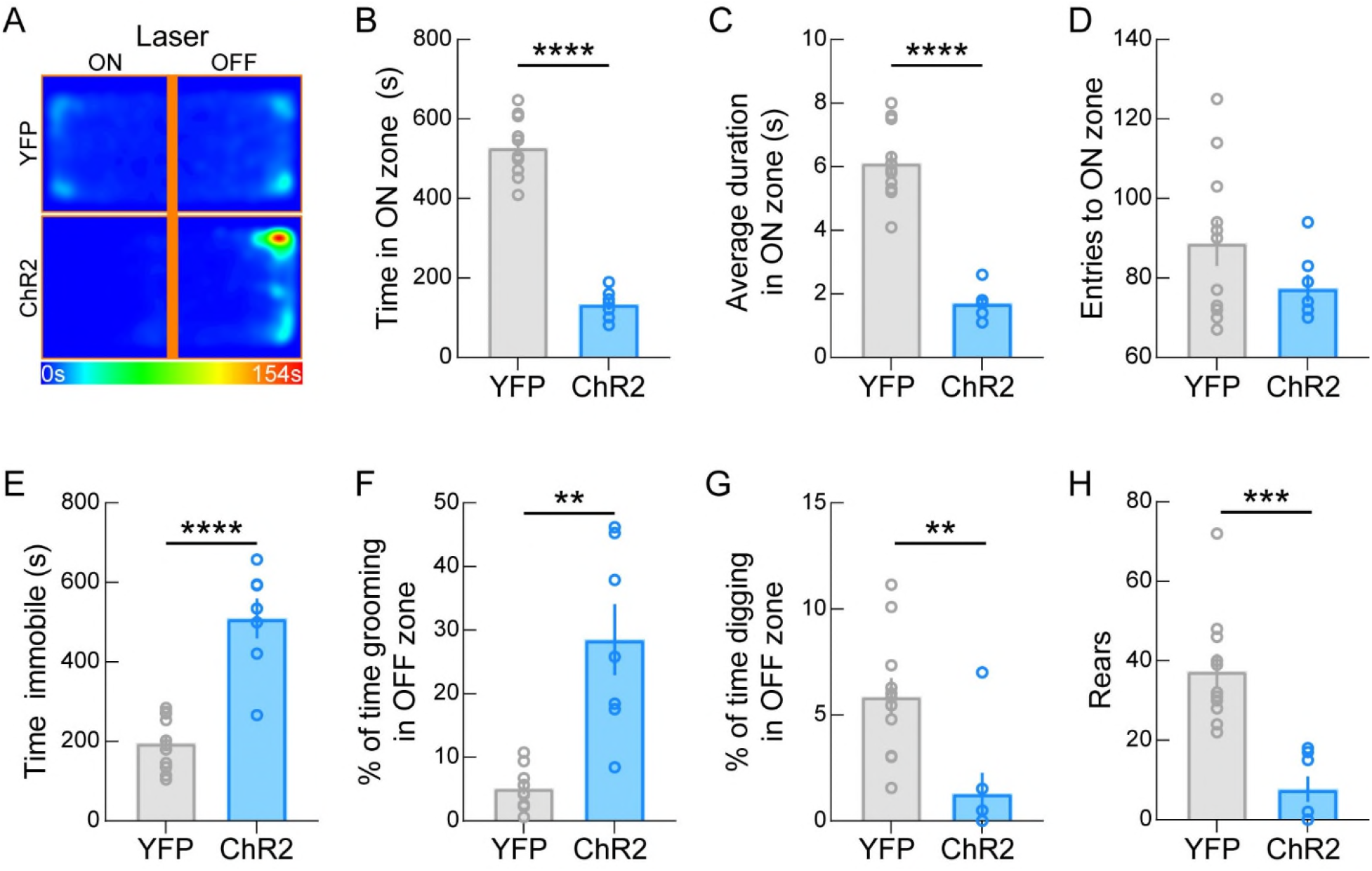
Photoactivation of the VMH→AHA pathway is aversive and triggers compulsive grooming behavior. **(A)** Mean heat maps for groups during RTPP show that ChR2 stimulation causes mice to spend more time in the Laser-OFF zone. **(B)** Time spent in the laser-paired ON zone is significantly reduced in ChR2-expressing mice compared to fluorophore controls. **(C)** The average duration of time spent in the Laser-ON zone is significantly reduced in ChR2-expressing mice compared to fluorophore controls. **(D)** There are no significant differences detected in the number of Laser-ON zone entries between groups. **(E)** Time immobile is significantly increased in ChR2 compared to fluorophore controls. **(F)** Time spent grooming in the Laser-OFF zone shows ChR2-expressing mice exhibit significantly more grooming behavior than fluorophore controls, even when normalized for time in the OFF zone. **(G)** VMH→AHA photostimulation results in a significant suppression of digging behavior in the Laser-OFF zone compared to fluorophore controls, even when normalized for time in the OFF zone. **(H)** A significant suppression of rearing behavior is observed in the ChR2-expressing mice compared to fluorophore controls. See **Supplementary Table 1** for corresponding statistics.

### Persistent VMH→AHA stimulation attenuates locomotion and promotes grooming over long timescales

The emergence of grooming behavior after exposure to predatory threat^24^ or environmental stress^25^ has been shown previously^9^. Thus, we wanted to test the effects of persistent stimulation of the VMH→AHA pathway on escape and post-escape behavior. For this, we used a fixed interval schedule for the delivery of light pulses consisting of 5 s “ON” and 25 s “OFF” for 3 hours on 3 consecutive days. Photostimulation evoked a significant increase in velocity on Day 1 (**Figure 4A**) but that effect was attenuated on subsequent days (**Figure 4B, C**) suggesting “deselection” of locomotion as escape behavior. Interestingly, across the 3 consecutive days, we observed a significant increase in grooming behavior during both photostimulation and post-photostimulation sessions (**Figure 4D**). There results demonstrate that persistent photostimulation of the VMH→AHA pathway evokes long-lasting significant increases in grooming behavior.

**Figure 4.**
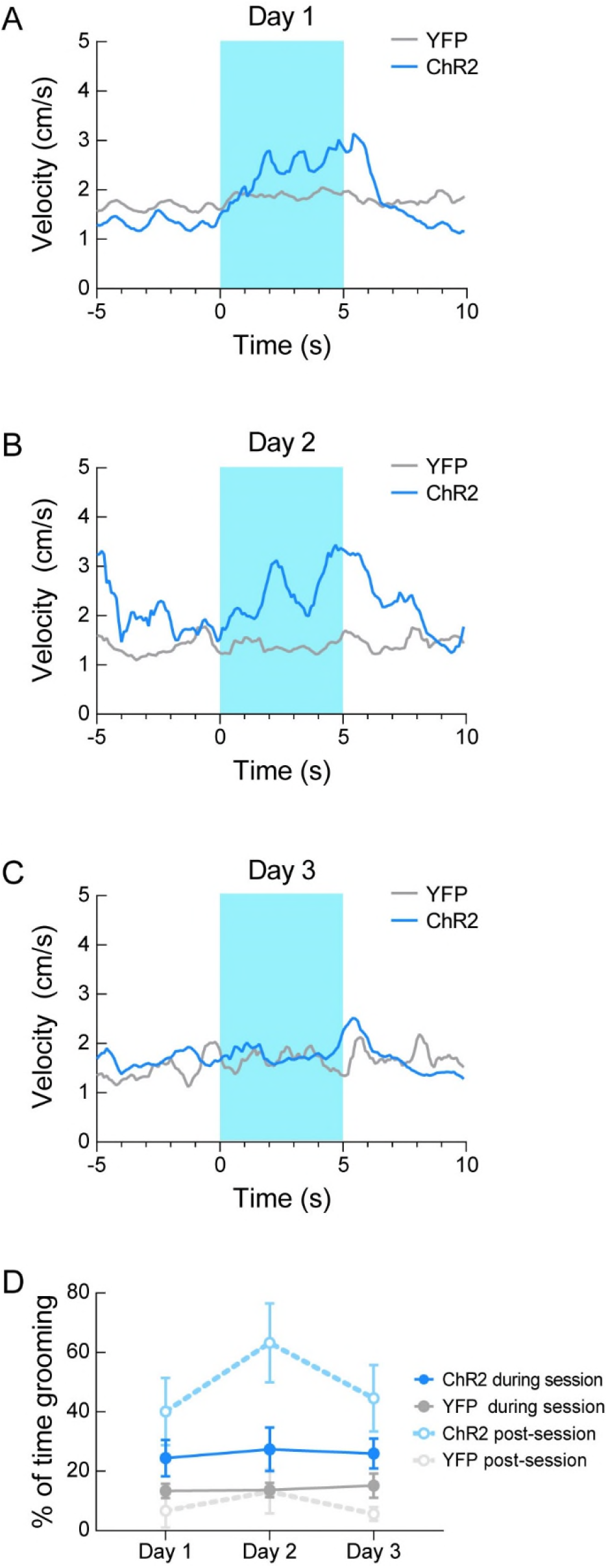
Persistent photoactivation of the VMH→AHA pathway attenuates locomotion and promotes long-lasting grooming behavior. **(A)** On Day 1, ChR2-expressing mice exhibit an increase in locomotor velocity upon photostimulation (mean of ∼360 trials per mouse, *n* = 4 mice). **(B)** On Day 2, ChR2-expressing mice exhibit an increase in locomotor velocity upon photostimulation that is indistinguishable from the pre-stimulation period (mean of ∼360 trials per mouse, *n* = 5 mice). **(C)** On Day 3, ChR2-expressing mice do not exhibit any change in locomotor velocity upon photostimulation (mean of ∼360 trials per mouse, *n* = 5 mice). **(D)** VMH→AHA stimulation increases the percent of time spent grooming post-session compared to fluorophore controls (*p* < 0.05 each day). Multiple comparisons reveal that grooming is greater in ChR2-expressing mice on Day 2 after the session (post) compared to behavior during the session. See **Supplementary Table 1** for corresponding statistics.

### Photoactivation of the VMH→AHA pathway increases neuronal activity in the amygdala

We conducted whole brain metabolic mapping using 2-deoxy-2-[18F]fluoro-D-glucose (FDG)-PET while photostimulating the VMH→AHA pathway in anesthetized mice. For this, we compared the effects of VMH→AHA activation in ChR2-expressing mice and control mice. We found that photostimulation of the VMH→AHA pathway significantly increased FDG uptake (i.e., increased metabolic activity) in brain areas encompassing the ventrolateral nucleus accumbens and the amygdala (**Figure 5**), which are brain region known to be involved in the regulation of fear-related behaviors^26-28^. Notably, VMH→AHA pathway activation was not associated with decreases in brain metabolic activity.

**Figure 5.**
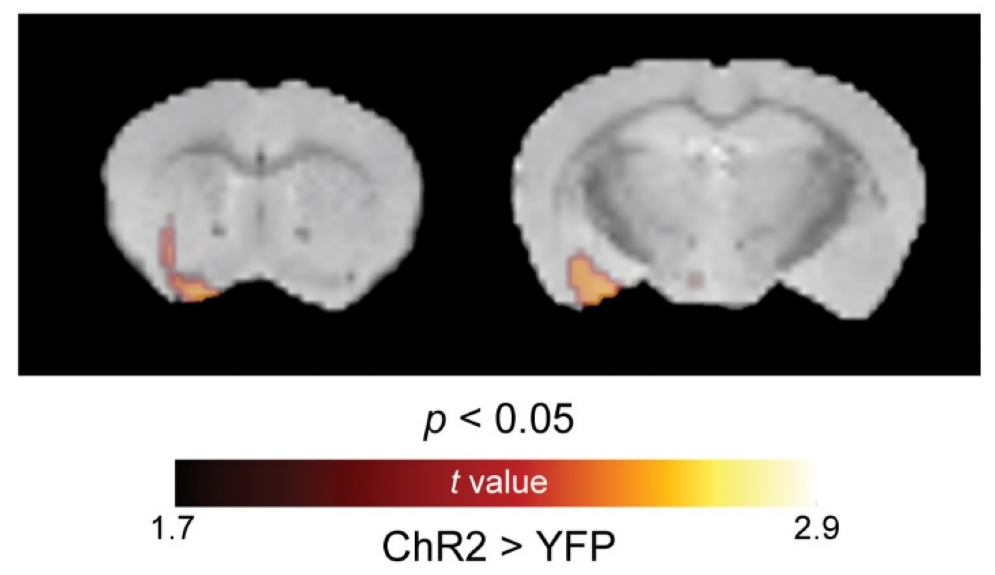
Photoactivation of the VMH →AHA pathway increases neuronal activity in the amygdala. FDG-PET overlaid on structural MRI shows that VMH→AHA stimulation significantly increases FDG uptake compared to controls (*p* < 0.05, unpaired *t*-test; *n* = 6 ChR2 and 10 YFP) in a region encompassing the extended amygdala.

## DISCUSSION

Fight-or-flight behavior requires mobilization of peripheral organ systems by the central nervous system (CNS) and is fundamental to fitness across mammalian species^29, 30^. Hypothalamic circuitry is involved in controlling defensive behaviors, particularly escape behaviors, through a network that involves the VMH, AHA, and dorsal premammillary nucleus (PMD)^7, 31^. Previous studies suggest that the VMH instantiates emotional states^32^. The AHA is a key structure in a dense interconnected hypothalamic network that regulates threat-induced behaviors^33^, and it exerts effects across physiological systems controlling thermoregulation^34^ and blood pressure^35^ that facilitate fight-or-flight behavioral selection^36^. Interestingly, GABAergic neurons in the AHA (AHA^VGAT^) promote biting behavior^37^. However, a study using the expression of FOS as an indirect marker for neuronal activity in the AHA revealed that fighting or mating did not evoke an increase in the number of FOS-positive neurons in the AHA^38^. Furthermore, electrolytic lesions of the VMH and AHA attenuate freezing in response to a predator odor^39^. Together, these studies further support a role of the AHA in defensive behaviors.

Here, we report that presentation of a foot-shock associated auditory cue evokes an increase in AHA neuronal activity demonstrating that AHA neurons are modulated by fear conditioning. Interestingly, on the day after shock-paired conditioning, we observed that AHA neuronal activity was significantly lower during the shock and post-shock periods established on Day 1 indicating temporal learning about the duration of the foot shock. Moreover, after shock-paired conditioning, significant freezing behavior emerges following the tone, further supporting learning within the group. While these experiments are insufficiently powered to identify sex differences, freezing behavior was exhibited by both sexes (females, *n* = 3 freezers and *n* = 1 non-freezer; males, *n* = 2 freezers and *n* = 2 non-freezers).

We used three behavioral assays to elucidate the effects on behavior during photostimulation of the VMH→AHA pathway. First, we performed an open field test as an unbiased assessment of behavior. Our results are consistent with previous findings^6^ as we observed significant effects on locomotor activity and jumping behavior evoked by VMH→AHA activation. Notably, a key advancement depicted by our work is that we found significant increases in grooming behavior between photostimulation bouts. Self-grooming behavior is considered to model components of Obsessive-Compulsive Disorder (OCD)^40^, and this self-grooming is comparable to the grooming behavior elicited by a foot-shock^41^. Moreover, previous studies showed that displacement grooming emerges during conflicts between two behavioral systems whereby low priority behavior, such as grooming, overrides high priority behavior, such as escape^42^. Mouse grooming does model etiological and symptomatic aspects of compulsive behaviors that translate to human OCD^40^. Mouse models of OCD, such as the SAPAP3 knock out mouse, exhibit increases in time spent grooming and number of grooming bouts^43^. Comparably, following escape behavior triggered by activation of single-minded 1 (SIM1)-expressing neurons in the paraventricular hypothalamus (PVH^SIM1^), mice exhibited increases in grooming behavior^44^. Importantly, the genetic complexity of OCD has led to challenges in biomarker identification^8^, which underscores the need to generate relevant models that link genetics, neuroanatomy, physiology, and behavior^45^. Adding weight to the considerable value of animal models is the evidence that selective serotonin and norepinephrine reuptake inhibitors responsible for reducing rodent-grooming behavior^46^ and normalizing cue-based reactivity^47^ also reduce repetitive behavior in humans^48^. Thus, it is possible that our results demonstrating that activation of the VMH→AHA pathway promotes grooming over long timescales could serve as a basis for future models to study neuropsychiatric disorders characterized by repetitive and compulsive behaviors.

Next, we used the RTPP paradigm to identify behaviors evoked by VMH→AHA activation. We found that the average duration to get to the photostimulation-paired side of the arena was significantly reduced while no differences were observed for the total number of entries. These results suggest that either the mice were still driven to explore the photostimulation-ON zone, or they needed more time to learn or consolidate fear conditioning. Nevertheless, the significant avoidance to stay on the photostimulation-paired side of the arena indicates that the predominant effect of VMH→AHA activation is redirection away from the photostimulation-ON zone. It is worthwhile to note the emergence of grooming behavior during the RTPP test. Previous studies showed that elevation in grooming behavior occurs at the expense of behaviors related to exploration of space such as rearing and digging^49^.

Additionally, we examined the effects of persistent stimulation of the VMH→AHA pathway on escape and post-escape behavior by using a fixed-interval stimulation paradigm. Remarkably, we found that persistent photostimulation across days promotes “deselection” of locomotion as an escape behavior but significantly increases grooming behavior during both photostimulation and post-photostimulation sessions. Our results not only extended findings on the context-dependent roles that hypothalamic circuits play on escape vigor^33^, but also demonstrate that persistent photostimulation of the VMH→AHA pathway evokes long-lasting grooming behavior. Changes in grooming behavior can be observed after rodents undergo stressful experiences^50^, and self-grooming behavior relates to the degree of stress in an inverted-U curve relationship^51^. Thus, in the absence of aversiveness, low self-grooming is observed, pre-exposure to stressful states increases grooming behavior, and highly aversive experiences occlude grooming ^52^. Importantly, in our experiments, grooming remained elevated following the three-hour session of persistent VMH→AHA stimulation. This suggests that the initial effect of stimulation may be adaptive to facilitate flight from an aversive stimulus followed by a shift in behavioral selection towards atypical grooming. It is noteworthy that other studies have found the emergence of grooming behavior after ChR2-mediated activation of escape^44^, which can be driven by the lateral hypothalamus (LH)^53^. However, our results are limited to mice that exhibited predominant hypothalamic viral transduction medial to the fornix which does not include the LH^54^.

The AHA projects to structures that are highly conserved within mammalian species, including the medial preoptic area, lateral hypothalamus, dorsomedial nucleus, capsule of the ventromedial nucleus, dorsal premammillary nuclei, and the central gray^55^. Notably, the AHA consists predominantly of GABAergic neurons as well as a population of thyroid hormone-dependent parvalbumin neurons^56^. The AHA receives projections from the lateral septum, paraventricular hypothalamus, ventromedial hypothalamus, paraventricular thalamus, and the ventral premammillary nucleus^37^. Therefore, the AHA circuitry conveys a complex array of synaptic inputs to modulate neuronal activity through downstream networks.

Using (PET) combined with optogenetic activation of the VMH→AHA pathway in anesthetized mice, we found that the amygdala is a downstream area activated by the stimulation of this pathway. The amygdala is a highly conserved structure involved in fear-related behaviors^57^. Previous studies showed that the central amygdala and bed nucleus of the stria terminalis serve as a subpallidal corridor that exhibits increased activity during exposure to aversive stimuli and uncertain/remote threat^58^. In mice, the activity of glutamatergic neurons in the amygdala increases in response to aversive stimuli. Moreover, self-grooming behavior is observed following stimulation of glutamatergic neurons in the amygdala^59^. Together, our findings and those of others suggest that neurons within the hypothalamic→amygdala circuitry play a role in regulating defensive behaviors. Our work highlights the need for future experiments to determine the specific neuron populations that modulate defensive-related behaviors. In addition, further investigations will be needed to examine VMH→AHA loss-of-function and other behavioral paradigms for context-dependent flexibility of behavioral selection.

In summary, our work has revealed that the AHA responds to threat and that such increases in activity are sufficient to trigger compulsive grooming behavior. Moreover, we demonstrated that persistent photostimulation of the VMH→AHA pathway evokes long-lasting grooming behavior. Furthermore, we showed that VMH→AHA stimulation increases activity in the amygdala. Thus, our findings support the continued investigations of the hypothalamic→amygdala circuitry in the regulation of defensive- and compulsive-related behaviors.

## ACKNOWLEDGEMENTS

The authors acknowledge with gratitude C. Lupica and G. Schoenbaum for discussions and comments on the manuscript, and S. Sarsfield for figure preparation and manuscript edits. Permission to publish miniscope drawing granted by Doric Lenses Inc. This work was supported by the National Institute on Drug Abuse Intramural Research Program (NIDA IRP) (ZIADA000595 and ZIADA000069), U.S. National Institutes of Health (NIH).

## AUTHOR CONTRIBUTIONS

Conceptualization B.T.L., M.S.A., and Y.A.; Data Curation B.T.L., M.S.A., A.J., L.J.E., O.S., and D.W.; Formal Analysis B.T.L., A.J., A.S.P., L.J.E., and O.S.; Writing – Original Draft B.T.L., M.S.A., and Y.A.; Writing – Review & Editing B.T.L., M.S.A., A.J., A.S.P., L.J.E., O.S., D.J.W., M.M., and Y.A.

