## Supplementary Figure 1 for "Activation of the fear-responsive anterior hypothalamic area promotes avoidance and triggers compulsive grooming behavior in mice"

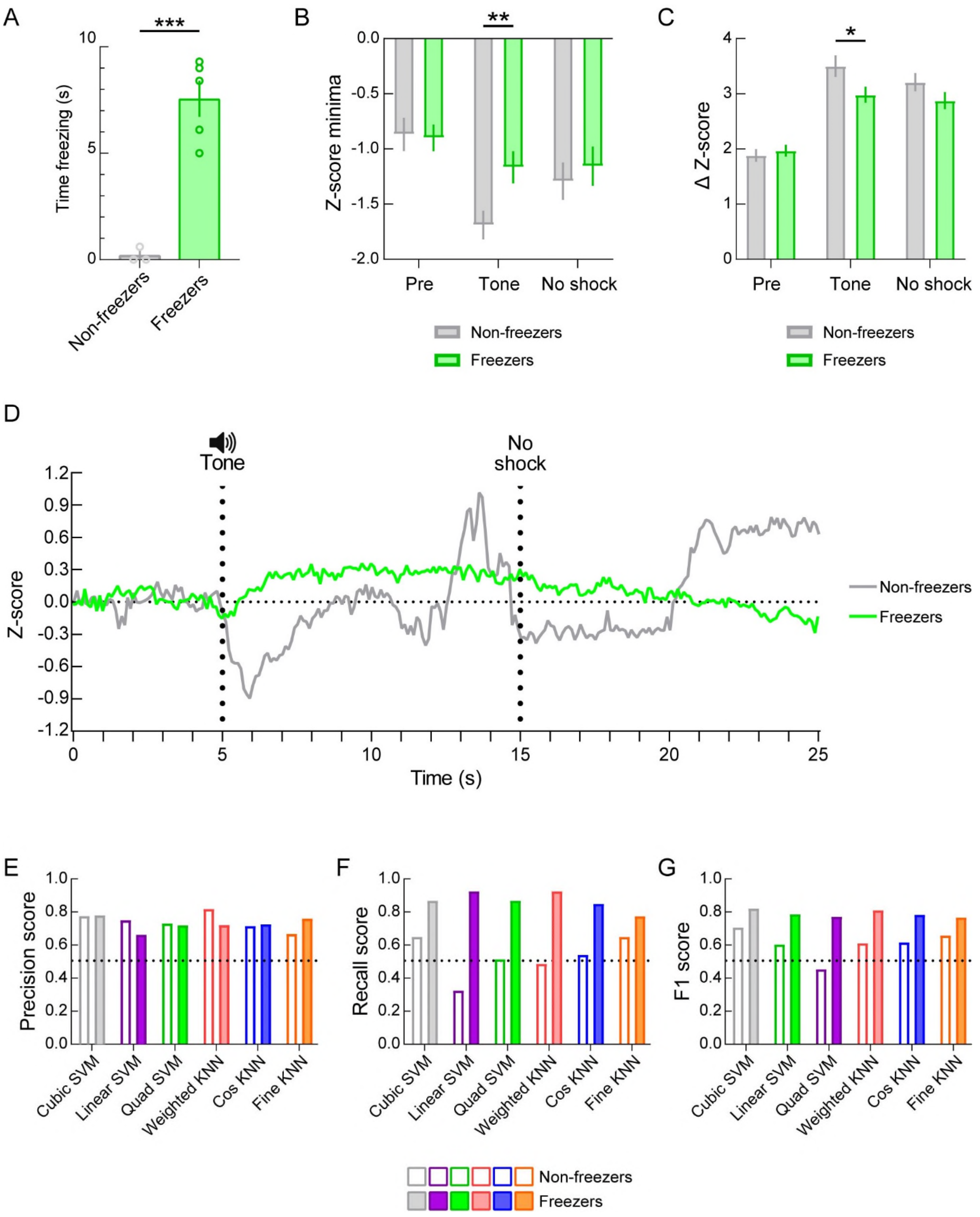

**Supplementary Figure 1. Machine learning model prediction of freezing versus non-freezing behavior after shock-cue conditioning.**

**(A)** Splitting the Day 2 trial 1 data into groups by whether mice exhibited freezing behavior (freezers) or not (non-freezers) reveals significant differences in time spent freezing. **(B)** Significantly lower Z-score minima are detected in non-freezers compared to freezers. **(C)** A significantly greater change in Z-score is detected in non-freezers compared to freezers during cue presentation. **(D)** Mean Z-score activity for all cells within each group shows a marked difference in AHA activity profiles. **(E)** Comparison of precision scores across machine learning models for classification of mice into freezer and non-freezer groups. All models perform above chance (dotted line) for precision. **(F)** Comparison of recall scores across machine learning models for classification of mice into their respective groups. All models except linear SVM and weighted KNN perform at or above chance (dotted line). **(G)** Comparison of F1 scores across machine learning models for classification of mice into their respective groups. Cubic SVM and fine KNN produce the best models. Quad SVM performed below chance (dotted line).
