## Supplementary Table 1 for "Activation of the fear-responsive anterior hypothalamic area promotes avoidance and triggers compulsive grooming behavior in mice"

**Supplementary Table 1. Statistical analyses**

| Figure | Test Name | Assumptions | Test Statistics | Test *p* value | Multiple Comparisons | *p* Value |
|---|---|---|---|---|---|---|
| Fig 1c | Welch's t-test; *n* = 8 animals (4 male, 4 female) | Variances significantly different, Welch's correction required | $t_{9.992}$ = 4.368 | $p = 0.0014$ | | |
| Fig 1i | Two way RM ANOVA, *n* = 120 cells from 8 animals day 1, 90 cells from 8 animals day 2 | Greenhouse-Geisser epsilon 0.66, sphericity not assumed, G-G correction applied | Stage = F(2.010, 418.0) = 71.37<br>Day = F(1, 216) = 76.19<br>Stage x Day = F(3, 624) = 82.20 | $p < 0.001$<br>$p < 0.001$<br>$p < 0.001$ | Within days<br>Day 1<br>Pre vs. Tone<br>Pre vs. Shock<br>Pre vs. Post<br><br>Day 2<br>Pre vs. Tone<br>Pre vs. Shock<br>Pre vs. Post<br><br>Between days<br>Pre<br>Tone<br>Shock<br>Post | <br><br>$p = 0.9948$<br>$p < 0.0001$<br>$p < 0.0001$<br><br><br>$p = 0.3315$<br>$p > 0.9999$<br>$p = 0.9742$<br><br><br>$p = 0.1798$<br>$p = 0.0046$<br>$p < 0.0001$<br>$p < 0.0001$ |
| Fig 1-1a | Welch's t-test, *n* = 5 freezers, 3 non-freezers | Variances nearly significantly different, Welch's correction applied | $t_{5.742}$ = 10.20 | $p < 0.0001$ | | |
| Fig 1-1b | 2-way RM ANOVA, *n* = 53 cells from 5 freezers, *n* = 37 cells from 3 freezers | Greenhouse-Geisser epsilon 0.7226, sphericity not assumed, | Cell: F(88, 176) = 2.766<br>Group: F(1, 88) = 1.544<br>Stage: F(1.445, 127.2) = 10.30<br>Stage x Group: F(2, 176) = 2.766 | $p < 0.0001$<br>$p < 0.2173$<br>$p < 0.0004$<br>$p < 0.0657$ | Within groups<br>Freezers<br>Pre vs. Tone<br>Pre vs. No shock<br><br>Non-freezers<br>Pre vs. Tone<br>Pre vs. No Shock<br><br>Between groups<br>Pre<br>Tone<br>No shock | <br><br>$p = 0.0349$<br>$p = 0.1969$<br><br><br>$p < 0.0001$<br>$p = 0.0832$<br><br><br>$p = 0.8723$<br>$p = 0.0084$<br>$p = 0.5831$ |

| | | | | | | |
|---|---|---|---|---|---|---|
| Fig 1-1c | 2-way RM ANOVA, *n* = 53 cells from 5 freezers, *n* = 37 cells from 3 freezers | Greenhouse-Geisser epsilon 0.8566, sphericity not assumed | Cell: F(88, 176) = 1.795<br>Group: F(1, 88) = 3.067<br>Stage: F(1.445, 127.2) = 56.28<br>Stage x Group: F(2, 176) = 2.689 | *p* = 0.0708<br>*p* < 0.0001<br>*p* = 0.0834<br>*p* = 0.0005 | Within groups<br>Freezers<br>Pre vs. Tone<br>Pre vs. No shock<br><br>Non-freezers<br>Pre vs. Tone<br>Pre vs. No Shock<br><br>Between groups<br>Pre<br>Tone<br>No shock | <br><br>*p* < 0.0001<br>*p* < 0.0001<br><br><br>*p* < 0.0001<br>*p* < 0.0001<br><br><br>*p* = 0.5867<br>*p* = 0.0357<br>*p* = 0.1428 |
| Fig 3a | Two-way RM ANOVA: Distance *n* = 5 ChR2, 11 YFP | Sphericity not assumed, matching is effective, mice 173987 and 173231 excluded for jump from apparatus | Stim x Transgene: F(5,70) = 1.907<br>Stim: F(3.594, 50.31) = 14.19<br>Transgene: F(1, 14) = 1.950<br>Subject: F(14, 70) = 9.117 | *p* = 0.1041<br>*p* < 0.0001<br>*p* = 0.1844<br>*p* < 0.0001 | Within groups<br>Between groups | ns<br>ns |
| Fig 3b | Two-way RM ANOVA: Jumps *n* = 7 ChR2, 11 YFP | Sphericity not assumed, matching is effective, | Stim x Transgene: F(5, 80) = 8.526<br>Stim: F(1.376, 22.02) = 8.526<br>Transgene: F(1,16) = 8.517<br>Subject: F(16, 80)= 3.337 | *p* < 0.0001<br>*p* = 0.0043<br>*p* = 0.0100<br>*p* = 0.0002 | Within groups<br>Between groups | ns<br>ns |
| Fig 3c | Two-way RM ANOVA: Time Immobile *n* = 5 ChR2, 11 YFP | Sphericity not assumed, matching is effective, mice 173987 and 173231 excluded for jump from apparatus impaired tracking, exclude first off bin | Stim: F(3.1, 43.39) = 2.861<br>Stim x Transgene: F(1, 16) = 4.792<br>Transgene: F(4,56) = 1.388 | *p* = 0.0312<br>*p* = 0.1415<br>*p* = 0.1513 | Within groups<br>Between groups | ns<br>ns |

| | | | | | | |
|---|---|---|---|---|---|---|
| Fig 3d | Two-way RM ANOVA: Grooming<br>*n* = 7 ChR2, 11 YFP | Sphericity not assumed, matching is effective, Sidak correction for multiple comparisons | Stim x Transgene: F(5, 80) = 5.027<br>Stim: F(2.509, 40.14) = 7.549<br>Transgene: F(1, 16) = 0.6117<br>Subject: F(16,80) = 5.188 | $p = 0.0005$<br>$p = 0.0008$<br>$p = 0.4456$<br>$p < 0.0001$ | Within groups<br>ChR2<br>Off1 vs. Off 2<br>On1 vs OfF1<br>All others<br>YFP<br>All<br><br>Between groups<br>Off 2<br>All others | <br><br>$p = 0.0097$<br>$p = 0.0213$<br>ns<br><br>ns<br><br><br>$p = 0.0253$<br>ns |
| Fig 3e | Two-way RM ANOVA: Rearing<br>*n* = 7 ChR2, 11 YFP | Sphericity not assumed, matching is effective, Sidak correction for multiple comparisons | Stim x Transgene: F(5, 80) = 2.648<br>Stim: F(2.858, 45.73) = 2.165<br>Transgene: F(1, 16) = 10.97<br>Subject: F(16, 80) = 3.976 | $p = 0.0288$<br>$p = 0.1079$<br>$p = 0.0044$<br>$p < 0.0001$ | Within groups<br>Between groups<br>Off3<br>On 3 | ns<br><br>$p = 0.0188$<br>$p = 0.0344$ |
| Fig 3f | Two way RM ANOVA: Time in center<br>*n* = 5 ChR2, 11 YFP | Sphericity not assumed, matching is effective, mice 173987 and 173231 excluded for jump from apparatus, exclude first off bin | Stim x Transgene: F(5, 70) = 2.409<br>Stim: F(3.260, 45.63) = 1.433<br>Transgene: F(1, 14) = 0.1650<br>Subject (14, 70) = 6.437 | $p = 0.0449$<br>$p = 0.2439$<br>$p = 0.6907$<br>$p < 0.001$ | Within groups<br>Between groups | ns<br>ns |
| Fig 4a | Welch's t-test: Time in on-zone<br>*n*= 7 ChR2; 11 YFP | F test to compare variance ($p = 0.0971$) | $t_{16} = 13.12$ | $p < 0.0001$ | | |
| Fig 4b | Welch's t-test: Avg duration of visit to on-zone<br>*n*= 7 ChR2; 11 YFP | F test to compare variance ($p = 0.0323$) | $t_{16} = 9.387$ | $p < 0.0001$ | | |
| Fig 4c | Welch's t-test: Entries to on zone<br>*n*= 7 ChR2; 11 YFP | F test to compare variance ($p = 0.0677$) | $t_{16} = 1.464$ | $p = 0.1627$ | | |

| | | | | | | |
|---|---|---|---|---|---|---|
| Fig 4e | Welch's t-test: Time immobile *n*= 7 ChR2; 11 YFP | F test to compare variance ($p = 0.0572$) | $t_{16} = 6.735$ | $p < 0.0001$ | | |
| Fig 4f | Welch's t-test: % time in off zone spent grooming *n*= 7 ChR2; 11 YFP | F test to compare variance ($p < 0.0001$) | $t_{6.317} = 4.130$ | $p < 0.0001$ | | |
| Fig 4g | Welch's t-test: number of rears *n* = 7 ChR2; 11 YFP | F test to compare variance ($p = 0.2161$) | $t_{16} = 4.969$ | $p < 0.0001$ | | |
| Fig 4h | Welch's t-test:% time in off zone spent digging *n* = 7 ChR2; 11 YFP | F test to compare variance ($p < 0.8103$) | $t_{14.05} = 3.497$ | $p < 0.0001$ | | |
| Fig 5a | Two-way RM ANOVA; *n* = 4 ChR2, 10 YFP; mice 173235, 173988, 173987 lost due to camera shift | Sphericity not assumed, matching is effective, some mice removed for acquisition tracking error | Stim x Transgene: F(1, 12) = 1.699<br>Stim: F(1, 12) = 6.592<br>Transgene: F(1, 12) = 3.274<br>Mouse: F(12, 12) = 1.205 | $p = 0.2169$<br>$p = 0.0247$<br>$p = 0.0955$<br>$p = 0.3760$ | Between Groups<br>Delta 1<br>Delta 2<br><br>Within Groups<br>ChR2<br>YFP | <br>$p = 0.0365$<br>$p = 0.6498$<br><br><br>$p = 0.0409$<br>$p = 0.2599$ |
| Fig 5b | Two-way RM ANOVA; *n* = 5 ChR2, 10 YFP; 173231 disconnected from laser; 173987 too blurry; 17988 door left open | Sphericity not assumed, matching is effective, some mice removed for acquisition tracking error | Stim x Transgene: F(1, 13) = 0.6551<br>Stim: F(1, 13) = 0.8470<br>Transgene: F(1, 13) = 0.1411<br>Mouse: F(13, 13) = 0.7684 | $p = 0.4329$<br>$p = 0.3742$<br>$p = 0.7132$<br>$p = 0.6791$ | Between Groups<br>Delta 1<br>Delta 2<br><br>Within Groups<br>ChR2<br>YFP | <br>$p = 0.6396$<br>$p = 0.9222$<br><br><br>$p = 0.5524$<br>$p = 0.9930$ |

| | | | | | | |
|---|---|---|---|---|---|---|
| Fig 5c | Two-way RM ANOVA;<br>$n$ = 5 ChR2, 10 YFP; scored same mice as day 2 | Sphericity not assumed, matching is effective, some mice removed for acquisition tracking error | Stim x Transgene: F(1, 11) = 0.001299<br>Stim: F(1, 11) = 1.165<br>Transgene: F(1, 11) = 0.2716<br>Mouse: F(11, 11) = 2.259 | $p$ = 0.9719<br>$p$ = 0.3034<br>$p$ = 0.6126<br>$p$ = 0.961 | Between Groups<br>Delta 1<br>Delta 2<br><br>Within Groups<br>Delta 1<br>Delta 2 | <br>$p$ = 0.6829<br>$p$ = 0.6544<br><br><br>$p$ = 0.4918<br>$p$ = 0.4181 |
| Fig 5d | Three-way RM ANOVA;<br>$n$ = 5 ChR2, 10 YFP | | Day: F(2, 82) = 1.838<br>Transgene: F(1, 82) = 5.517<br>Phase: F( 1, 82) = 48.27<br>Day x Transgene: F(2, 82) = 1.377<br>Day x Phase: F(2,82) = 0.6065<br>Transgene x Phase: F(1,82) = 14.59<br>Day x Transgene x State: F(2,82) = 0.3039 | $p$ = 0.1656<br>$p$ = 0.212<br>$p$ < 0.0001<br>$p$ = .2582<br>$p$ = 0.5477<br>$p$ = 0.0003<br>$p$ = 0.7388 | Within groups<br>ChR2<br>Day 1 In vs Post<br>Day 2 In vs Post<br>Day 3 In vs Post<br><br>YFP<br>Day 1 In vs Post<br>Day 2 In vs Post<br>Day 3 In vs Post<br><br>Between groups<br>Day 1 In<br>Day 2 In<br>Day 3 In<br><br><br>Day 1 Post<br>Day 2 Post<br>Day 3 Post | <br><br>ns<br>$p$ = 0.0471<br>ns<br><br><br>ns<br>ns<br>ns<br><br><br>ns<br>ns<br>ns<br><br><br>$p$ = 0.0151<br>$p$ < 0.0001<br>$p$ < 0.0055 |
| Fig 6a | T-test with probabilistic Threshold-free Cluster Enhancement (pTFCA) | | Heatmap for $t$ scale on figure | $p$ < 0.05 | | |

| | | | | | | |
|---|---|---|---|---|---|---|
| SFig 1a | Welch's t-test: Avg duration of visit to on-zone<br>*n*= 3 Non-Freezers; 5 Freezers | Due to small 'n' and nearly significant differences of variances, Welch's correction applied, one 'Freezer' mouse excluded due to error in miniscope analysis | $t_{4.427} = 8.417$ | $p = 0.0007$ | | |
| SFig 1b | Two-way RM ANOVA with uncorrected fishers LSD | Data approximately normal, Sphericity not assumed, data from one 'freezer' mouse excluded | Stage x Group: F(2, 176) = 2.766<br>Stage: F(1.445, 127.2) = 10.30<br>Group: F(1, 88) = 1.544<br>Cell: F(88, 176) = 2.90 | $p = 0.0657$<br>$p = 0.0004$<br>$p = 0.2173$<br>$p < 0.0001$ | Within Groups<br>Freezers<br>Pre vs. Tone<br>Pre vs. No shock<br>No Freezers<br>Pre vs. Tone<br>Pre vs. No shock<br><br>Between Groups<br>Pre<br>Tone<br>No Shock | <br><br>$p = 0.0349$<br>$p = 0.1969$<br><br>$p < 0.0001$<br>$p < 0.0832$<br><br><br>$p = 0.8723$<br>$p = 0.0084$<br>$p = 0.5831$ |
| SFig 1c | Two-way RM ANOVA with uncorrected fishers LSD | Data approximately normal, Sphericity not assumed, data from one 'freezer' mouse excluded | Stage x Group: F(2, 176) = 2.689<br>Stage: F(1.445, 127.2) = 56.28<br>Group: F(1, 88) = 3.067<br>Cell: F(88, 176) = 1.795 | $p = 0.0708$<br>$p < 0.0001$<br>$p = 0.0834$<br>$p = 0.0005$ | Within Groups<br>Freezers<br>Pre vs. Tone<br>Pre vs. No shock<br>No Freezers<br>Pre vs. Tone<br>Pre vs. No shock<br><br>Between Groups<br>Pre<br>Tone<br>No Shock | <br><br>$p < 0.0001$<br>$p < 0.0001$<br><br>$p < 0.0001$<br>$p < 0.0832$<br><br><br>$p = 0.5867$<br>$p = 0.0357$<br>$p = 0.1428$ |
